# Medial prefrontal cortex represents the object-based cognitive map when remembering an egocentric target location

**DOI:** 10.1101/680199

**Authors:** Bo Zhang, Yuji Naya

## Abstract

A cognitive map, representing an environment around oneself, is necessary for spatial navigation. However, compared with its constituent elements such as individual landmarks, neural substrates of coherent spatial information remain largely unknown. The present study investigated how the brain codes map-like representations in a virtual environment specified by the relative positions of three objects. Representational similarity analysis revealed an object-based spatial representation in the hippocampus (HPC) when participants located themselves within the environment, while the medial prefrontal cortex (mPFC) represented it when they recollected a target object’s location relative to their self-body. During recollection, task-dependent functional connectivity increased between the two areas implying exchange of self- and target-location signals between the HPC and mPFC. Together, the coherent cognitive map, which could be formed by objects, may be recruited in the HPC and mPFC for complementary functions during navigation, which may generalize to other aspects of cognition, such as navigating social interactions.

## Introduction

During navigation, it is necessary to locate our self-position in the current spatial environment as well as to locate the objects relative to the self-body (i.e., egocentric location). To conduct each of the two mental operations, we need map-like representations, called “cognitive map” in our brain (Tolman, 1948). After the discovery of “place cells,” the hippocampus (HPC) of the medial temporal lobe (MTL) has been considered responsible for the cognitive map (Buffalo, 2015), and crucial contributions of the HPC to spatial memory have also been reported by animal model studies that evaluated behavioral patterns of rodents with an inactivated HPC using maze tasks (Nakazawa et al., 2002; Packard and McGaugh, 1996; Redish and Touretzky, 1998) as well as human studies that demonstrated the relationship between HPC volume in individual subjects and their amounts of experience exploring spatial environments (Woollett and Maguire, 2011; Schinazi et al., 2013, e.g., London taxi drivers). However, it remains largely unknown how neural substrates of the cognitive map are involved in the two mental operations required to locate specific objects within the environment. One possible reason for the difficulty in addressing this question is that despite extensive studies on the spatial elements related to the cognitive map (e.g., self-location, head-direction etc.) (O’Keefe and Dostrovsky, 1971; Vass and Epstein, 2013; Chadwick et al., 2015; Buffalo, 2015; McCormick et al., 2018;), there is still a lack of sufficient isolation and characterization of the neural signal of the cognitive map under the previous research paradigms.

In addition to the HPC, the role of the medial prefrontal cortex (mPFC) in goal-directed planning during navigation was demonstrated by a previous human fMRI study that showed increased connectivity between the HPC and mPFC (Brown et al., 2016). The mPFC has been long considered as a member of the core-brain system in the retrieval of episodic memory (Konishi et al., 2000; Eichenbaum, 2017; McCormick et al., 2018), which is an autobiographical memory consisting of spatial, object, and temporal information (Suzuki and Naya, 2011; Naya and Suzuki, 2011; Squire and Wixted, 2011). Schacter et al. (2007) suggested an involvement of the mPFC in future-simulation processing and recollection of past episodes, which depend on mnemonic information stored as declarative memory including both episodic and semantic memory. Recently, they also showed increased connectivity between the HPC and mPFC during future simulation (Campbell et al., 2018). This preceding literature suggests that the HPC and mPFC, which belong to the default-mode network, work together when remembering stored information (e.g., cognitive map) and construct the mental representation of goal-directed information (e.g., target-location) from mnemonic information with the current context (e.g., self-location) (Schacter, 2012). However, the specific functional role of each of HPC and mPFC during the construction process (McCormick et al., 2018; Campbell et al., 2018) remain elusive, presumably because the construction of goal-directed information (e.g., spatial navigation) includes at least two mental operations described above (locating the self and locating an object target relative to self-location), as previous experimental paradigms did not dissociate these aspects of behavior.

To address these issues, we aimed to devise a novel 3D spatial-memory task with spatial environments defined by objects, which would enable us to identify the representation of the cognitive map and to investigate how it is related to the two mental operations (Fig. 1). We used a stimulus set consisting of three different human characters throughout the entire experiment, while the spatial configuration of the three characters was changed in a trial-by-trial manner. The spatial configuration pattern was referred to as a *“map”* in the present study (Fig. 1b). In each trial, participants encoded a *map* from the first-person’s view by walking toward the characters in one of four fixed walking directions (walking period, Fig. 1c, see Methods). Following the walking period, one human character (facing object) was presented on a virtual-environment background with other characters being invisible, which gave the participants a feeling of facing the presented character in the virtual environment (i.e., facing period). After a short delay, one of the two remaining characters (targeting object) was presented without the virtual environment background, and the participants were required to remember the location of this second human character relative to their self-body (i.e., targeting period). Thus, the two mental operations were separated into two periods within a single trial. This task design allowed us to detect the brain regions that distinguished the spatial configurations of the objects (i.e., *map*) around the participants during the facing period and targeting period separately. The results of the representational similarity analysis (RSA; see Methods for details) (Kriegeskorte et al., 2012; Kriegeskorte et al., 2008) showed that the spatial environment defined by the three objects were represented in the HPC during the facing period, while it was represented in the mPFC during the targeting period, suggesting different contributions of the object-based cognitive map to the recollection between the two brain areas of the default-mode network.

**Figure 1:**
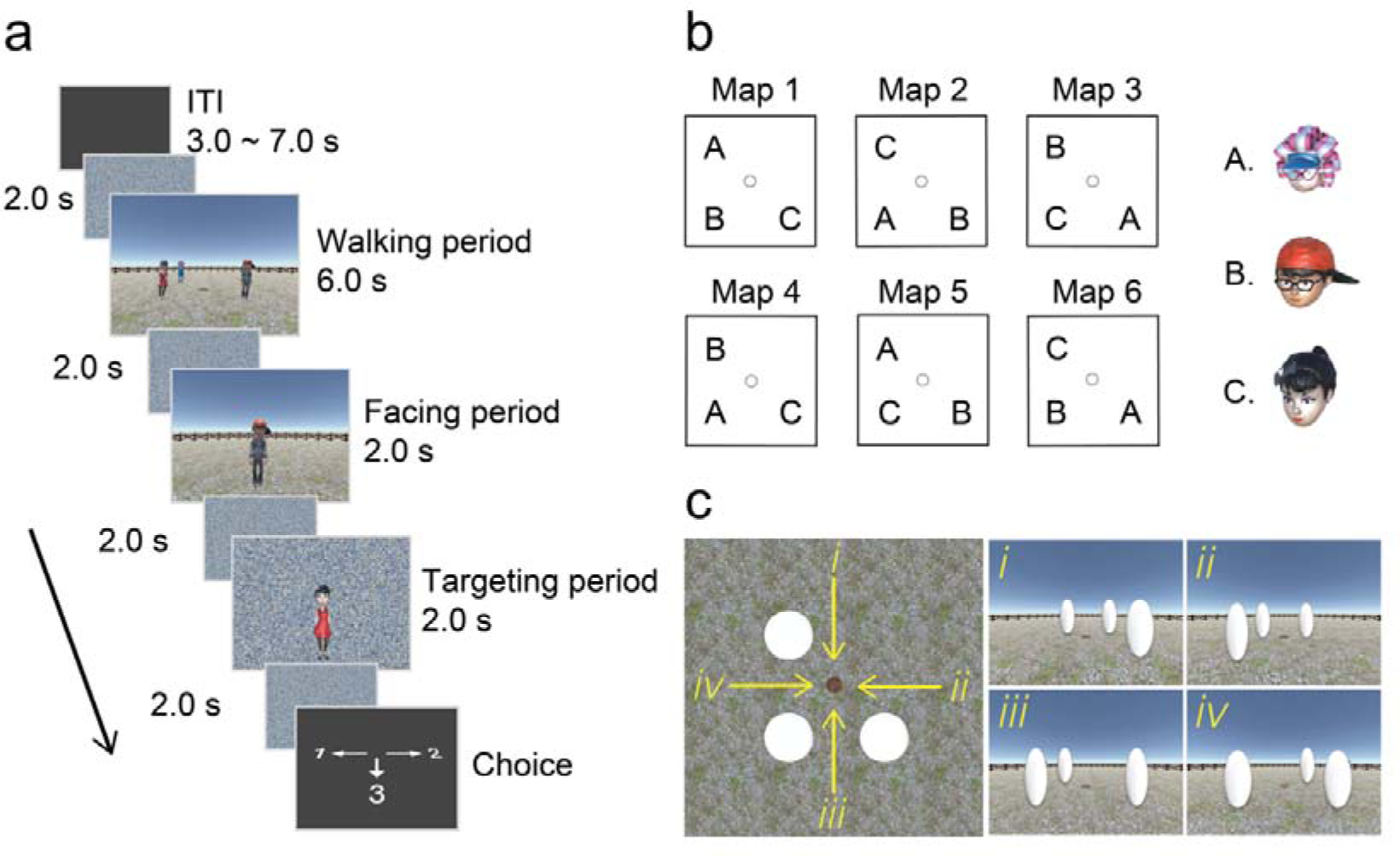
Task design. (a) spatial-memory task. Each trial consisted of three periods. In the walking period, participants walked toward three human characters using the first-person perspective and stopped on a wood plate. In the facing period, one of the human characters was presented, indicating the participant’s current self-orientation. In the targeting period, a photo of another character was presented on the scrambled background. The participants chose the direction of the target character relative to their body upon presentation of a response cue. (b) Maps were defined by the relative position of the three human characters, while the unfilled dot represents the wood plate. (c) The walking directions were defined by the spatial layout of the three human characters from the participant’s first-person perspective.

## Results

The experiment was conducted over two days with 19 participants. On the first day, the participants were familiarized with the 3D virtual environment and the three human characters through a head-nodding detection (HND) task (Fig. S1a). In this task, the participants had the same walking experience as in the spatial-memory task but were subsequently asked to indicate whether one of the three characters in a photo had nodded its head during the walking period. On the second day, the participants performed the spatial-memory task during fMRI scanning (Fig. 1a). To prevent voluntary memorization of the spatial relationship of the human characters during the walking period, the HND trials were pseudo-randomly mixed with the spatial-memory trials, and the participants were instructed to focus on head-nodding of the human characters during the walking period in all trials. In each trial, its trial-type (i.e., HND or memory task) became distinguishable only after the walking period by subsequent stimuli. All participants exhibited ceiling performance with a 93.6% ± 0.02% correct rate (mean ± SE, n = 19) for the spatial-memory task and no significant difference was found among each of the task parameters (e.g., maps, walking directions) (Fig. S1c). All participants also showed accuracy that was significantly higher than chance level (50%) in both the head-nodding and no head-nodding trials in the HND task (Fig. S1a). Attempts to memorize the spatial arrangements of the human characters during scanning were examined using post-scanning questionnaires. All participants reported that they did not make any voluntary effort to memorize the spatial relationship of the three human characters nor utilize any special strategy for memorizing it (Table S2). It should be noted that no participant was able to recall the number of map patterns they experienced in the experiment even though only three of the six possible patterns of maps were repeatedly presented to each participant. In addition, no significant changes in performance was found across four experimental sessions (Fig. S1b; F(3,72) = 0.38, P = 0.76), suggesting that the participants performed the spatial-memory task with high-performance from the beginning and did not learn to use a systematic strategy to improve their performance during the sessions. These behavioral results suggest that the present experimental design allowed us to investigate neural operations for the retrieval after the participants automatically encoded the spatial configuration of three human characters during the walking period when viewing the characters attentively to detect head-nodding.

## Neural representation of the object-based cognitive map

To decode the map information across the whole brain, we conducted searchlight-based RSA, which examined the multi-voxel pattern similarity between trial pairs in the “same map” and compared it with that in the “different map” condition across each brain voxel by drawing a 6 mm radius sphere with each voxel in the spherical center. Map information was decoded regardless of other task parameters such as the walking direction or the identity of the facing character by balancing the number of trials with other task parameters across the same and different map conditions in the experimental design as well as excluding the effects of other task parameters in the regression analysis (see Methods for details, Table. S1 for the regressor list and the averaged r^2^ among the regressors in each GLM).

We first assessed the map representation during the facing period (4.0 s including the subsequent delay; Fig. 2a), in which the participants oriented themselves to a presented human character in the 3D environment. We found a cluster located in the left middle HPC (mHPC; Fig. 2b, P < 0.01, voxel-wise threshold; P < 0.05, cluster-corrected for multiple comparison), suggesting that the map defined by multiple objects is represented in the HPC. In addition to the mHPC, clusters were revealed in the insula, angular gyrus, and superior temporal cortex (Fig. S2b, P < 0.01, voxel-wise threshold; P < 0.05, cluster-corrected for multiple comparison; see discussion). We next assessed the map representation during the targeting facing period (4.0 s including the subsequent delay), in which the participants remembered the location of a target character relative to their self-body (egocentric target location). In contrast to the facing period, we found clusters representing the map information mainly in the mPFC including the rectus and BA 10 (Fig. 2c) rather than in the HPC during the targeting period. A peak was revealed in the rectus (x = 4, y = 50, z = −18; t value: 5.62). Outside of the mPFC, we found clusters in the precuneus and middle temporal gyrus, and the inferior frontal cortex (Fig. 2c, Fig. S2c; P < 0.001, voxel-wise threshold; P < 0.05, cluster-corrected for multiple comparison). These three brain regions have been consistently reported to be involved in scene construction during recalling of past experience and imagination of new experiences (Hassabis and Maguire, 2007; Bird et al., 2010; Gaesser et al., 2013), which is consistent with the post-scanning report that all participants recalled and also imagined the egocentric positions of the three human characters during the targeting period.

**Figure 2:**
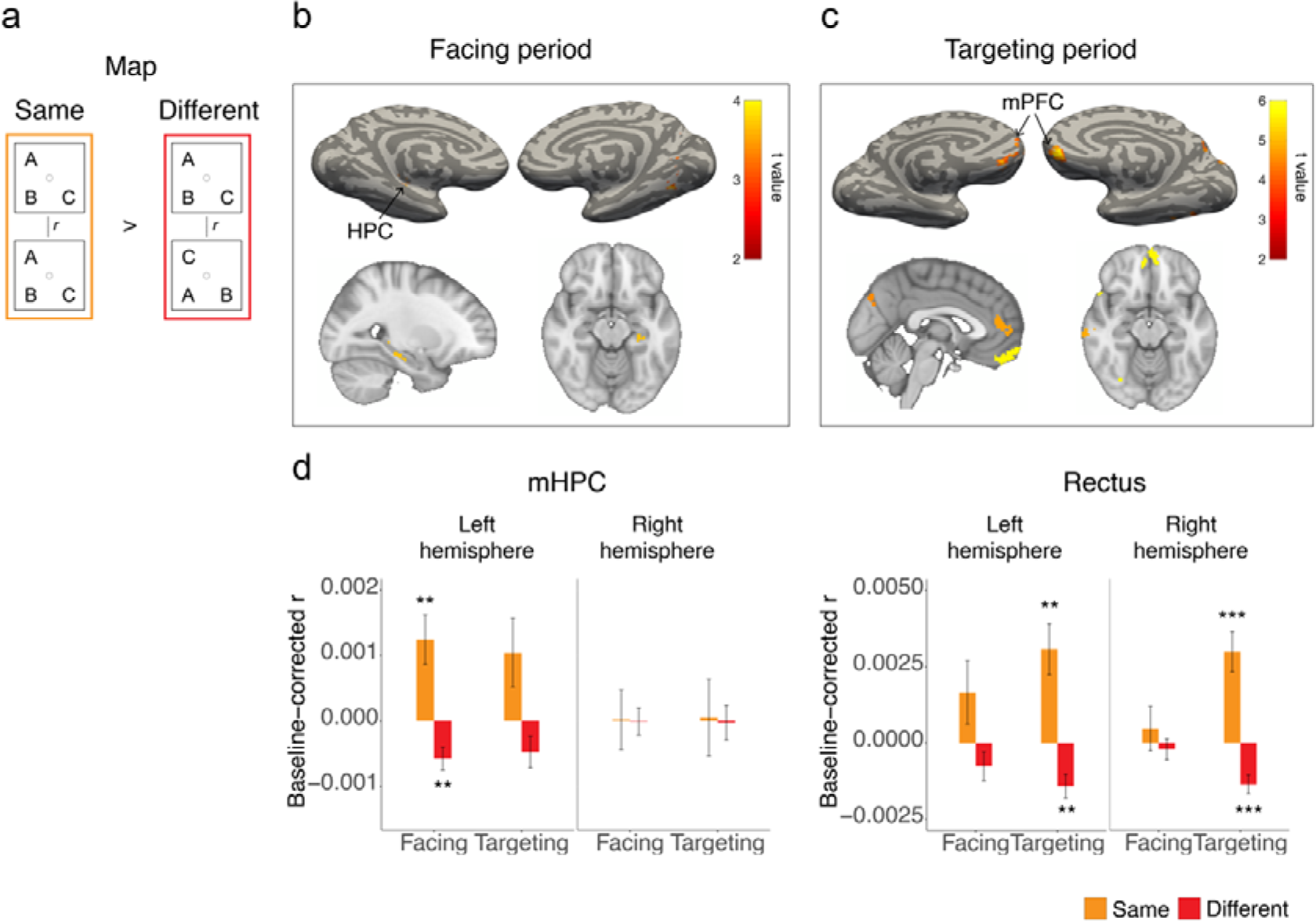
Neural representation of cognitive map in MTL and mPFC. (a) Schematic representation of decoding the maps using RSA. (b) In the facing period, RSA revealed a cluster in the left middle HPC (mHPC; t value: 3.84; MNI coordinates: −28, −25,−16; shown on sagittal and transverse section) within the MTL (P < 0.01, initial threshold; P < 0.05, cluster-corrected for multiple comparison). (c) In the targeting period, clusters were revealed in the mPFC (P < 0.001, initial threshold; P < 0.05, cluster-corrected for multiple comparison). The peak was revealed in the rectus within the mPFC (t value: 5.62; MNI coordinates: 4, 50, −18). (d) The left mHPC and bilateral rectus revealed a significantly higher similarity than chance level in the “same-map” condition during facing and targeting period, respectively (the left mHPC: t(18) = 3.26, P = 0.016; the left rectus: t(18) = 3.68, P = 0.007; the right rectus: t(18) = 4.50, P = 0.001; bonferroni corrected for multiple comparisons).

To validate the searchlight-based RSA results showing the map representations in the HPC and mPFC, we conducted a region of interest (ROI)-based RSA using a total of four masks consisting of both right and left hemispheres of the mHPC and rectus (mPFC) that derived from the automated anatomical labeling (AAL, Fig. S4a, bottom panel) template. In each ROI, we examined the multi-voxel pattern similarity in the “same-map” condition and in the “different-map” condition separately. These similarities were then compared with chance levels that were estimated as mean values among 5,000 of trial-based permutation results (see Methods).

In the left mHPC, the similarity was significantly higher than the chance level in the same-map condition during the facing period (Fig. 2d; t(18) = 3.26, P = 0.016, bonferroni corrected for multiple comparisons (n = 4)), while it was significantly lower than the chance level in the different-map condition (t(18) = −3.26, P = 0.016). This tendency was also observed during the targeting period, although the similarities for neither of the two conditions reached a statistical significance (same-map: t(18) = 1.99, P = 0.062; different-map: t(18) = −1.99, P = 0.062). In contrast to the left hemisphere, the similarities did not differ from the control levels in any combination of the conditions and periods in the right mHPC (Fig. 2d), suggesting a striking laterality effect on the map representation in the HPC. On the other hand, both hemispheres of the rectus showed significantly higher and lower similarities than the control levels in the same-map (the left rectus: t(18) = 3.68, P = 0.007; the right rectus: t(18) = 4.50, P = 0.001) and different map conditions (the left rectus: t(18) = −3.68, P = 0.007; the right rectus: t(18) = −4.49, P = 0.001), respectively, only during the targeting period. This tendency was observed during the facing period only in the left hemisphere, although the similarities for neither of the two conditions reached a statistical significance (same-map: t(18) = 1.60, P = 0.126; different-map: t(18) = −1.60, P = 0.126). The ROI-based RSA thus confirmed the results of the searchlight-based RSA, indicating that while the cognitive map was represented in the HPC during the facing period, the mPFC represented it during the targeting period. It should be noted here that the map representation in the different brain areas (i.e., HPC and mPFC) during the facing and targeting periods could not be explained by different background images during the two periods themselves (i.e., environment vs. noise) because the RSA revealed brain regions that discriminated trials with different map information during each period in which the same background was presented. Taken together, the HPC and mPFC may represent the cognitive map information for the different functional operations. Some temporally-overlapping map representation in the two brain areas may implicate an interaction to share the map information between them although it did not reach a statistical significance (i.e., map representation in the mPFC and HPC during the facing and targeting periods, respectively).

## Input signal for the map construction

To examine possible signal input from the MTL sub-regions to the left mHPC for the map construction during the facing period, we examined the neural representation of the facing-character identity and walking direction that the participants perceptually and/or mentally re-experienced based on their post-scanning reports (Table S2; Fig. S3a). The results revealed that the bilateral perirhinal cortex (PRC) encoded character identity (Fig. S3b; P < 0.001, initial threshold; P < 0.05, cluster-corrected for multiple comparison) (Naya et al., 2001; Suzuki and Naya, 2014), while the parahippocampal cortex (PHC) and left retrosplenial cortex (RSC) encoded the walking directions reflecting the spatial layout of one empty and three occupied positions perceived by the participants during the walking period (Fig. 1c). In the HPC, the left posterior HPC (pHPC) selectively represented the spatial layout but not the character identity, while the bilateral anterior HPC (aHPC) revealed clusters for both character identity and spatial layout (Fig. S3b). These results were consistent with the notion of the “two cortical systems” model suggesting that object identity and spatiotemporal context are processed in two separate neural systems with the PRC and PHC-RSC as the core brain regions, with the two different information domains interacting in the HPC (Ranganath and Ritchey., 2012). Together, the RSA analyses suggest that the MTL is associated with representing the spatial environment in the following ways: elements such as each object identity and spatial layout are represented by extrahippocampal areas while the relative relationship between multi-objects is represented in the HPC, suggesting cognitive map representation in the HPC.

## Current self-orientation on the map

To compute the egocentric location of a target object (e.g., left, right, or back), information on the current self-position/orientation on the map is necessary (Fig. 3a). Therefore, we examined which brain regions were involved in representing such allocentric “heading-direction” signals (Hargreaves et al., 2005; Wang et al., 2018). Interestingly, while no significant cluster was revealed during the facing period (facing character; Fig. 3b), robust clusters were revealed during the targeting period (Fig. 3b, P < 0.001, voxel-wise threshold; P < 0.05, cluster-corrected for multiple comparison). These clusters were located in the left ERC, bilateral HPC and PHC inside the MTL as well as in the lateral occipital cortex, parietal cortex, precuneus, and anterior cingulate cortex outside the MTL (Fig. 3b, P < 0.001, voxel-wise threshold; P < 0.05, cluster-corrected for multiple comparison). These results suggested that a self-orientation signal was induced during the targeting period, presumably because of the necessity to compute the egocentric target location. This interpretation is consistent with the post-scanning report in which participants reported imagining their self-orientation on the map only during the targeting period.

**Figure 3:**
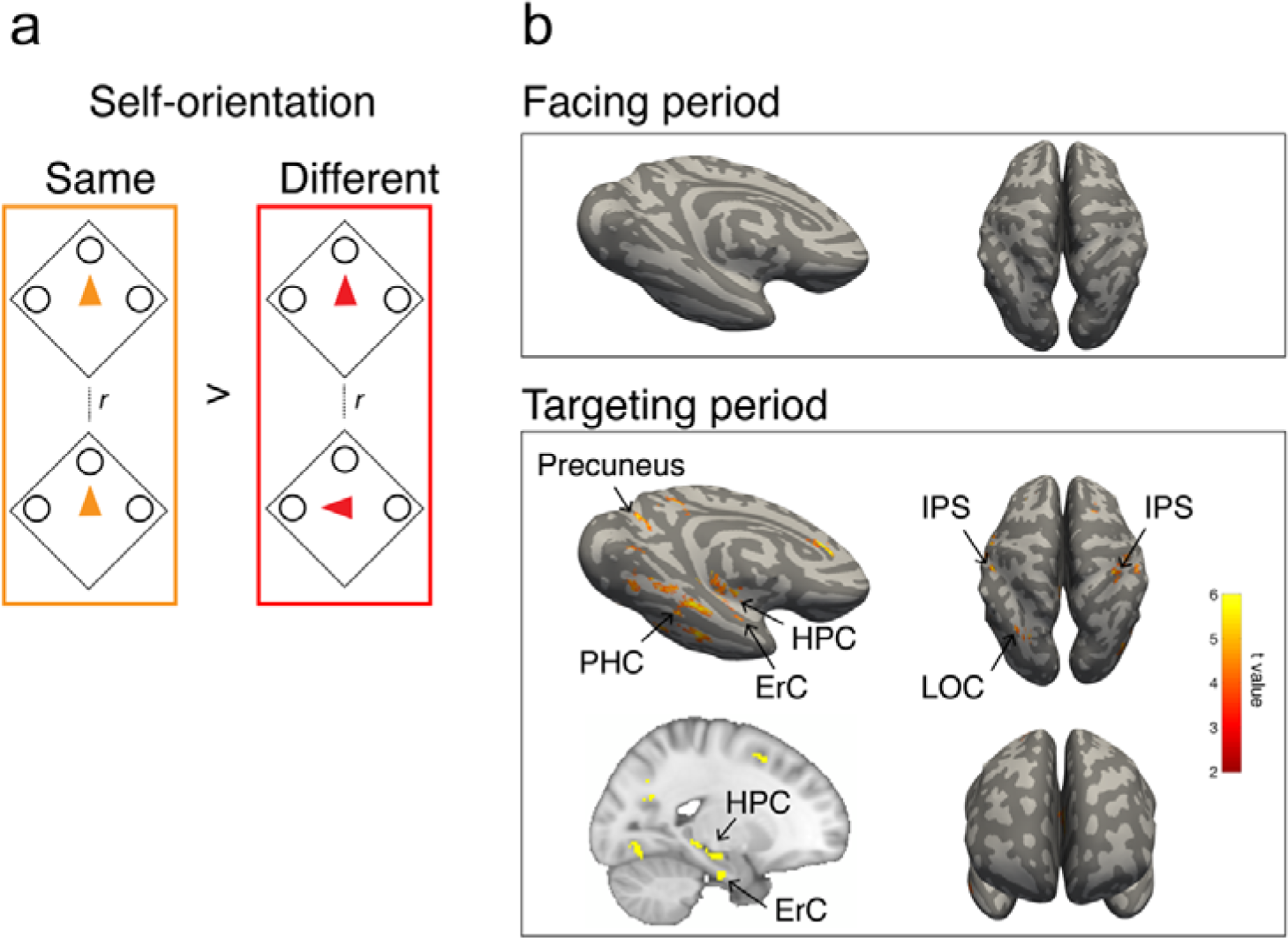
Neural representation of self-orientation on cognitive map. (a) Schematic representation of decoding participants’ self-orientation. (b) In the facing period, no cluster was revealed even with the use of a more liberal threshold (P < 0.01, initial threshold; P < 0.05, cluster-corrected for multiple comparison). In the targeting period, clusters were revealed in the MTL (bilateral HPC, PHC, and left ErC) and self-motion areas (inferior parietal cortex, RSC, and lateral occipital cortex).

## Remembering the egocentric location of a target object

Next, we examined which brain regions signaled the egocentric location (left, right, or back relative to self-body) of a target object (Fig. 4a). The results revealed robust clusters in both the mPFC and MTL (Fig. 4a, P < 0.001, voxel-wise threshold; P < 0.05, cluster-corrected for multiple comparison). In the mPFC, we identified the rectus, medial/superior orbitofrontal cortex, and olfactory cortex. In the MTL, clusters were found in the anterior HPC. Apart from the mPFC and MTL, clusters were also found in the lateral occipital cortex, inferior parietal cortex, anterior temporal lobe, premotor cortex, and lPFC (middle and superior PFC). We also found clusters in the precuneus and posterior parietal cortex, which were previously reported to represent the egocentric location (Chadwick et al., 2015). The widely distributed clusters may indicate that the brain regions representing the egocentric target locations can be involved in either generation of the egocentric-target-location information from multiple pieces of information (cognitive map, self-orientation, and target character identity) or its maintenance while preparing for the following response. These distinct functions might be supported by three different large-scale brain networks: the dorsal attention network, frontoparietal control network, and default-mode network (Spreng and Schacter., 2011). In contrast to the robust signal observed across different brain networks for egocentric target-location, no cluster was revealed for allocentric target location relative to the spatial layout of the characters (Fig. 4b, P < 0.001, voxel-wise threshold; P < 0.05, cluster-corrected for multiple comparison), which implies that the target location may be directly retrieved in the form of egocentric coordinates rather than via its allocentric representation.

**Figure 4:**
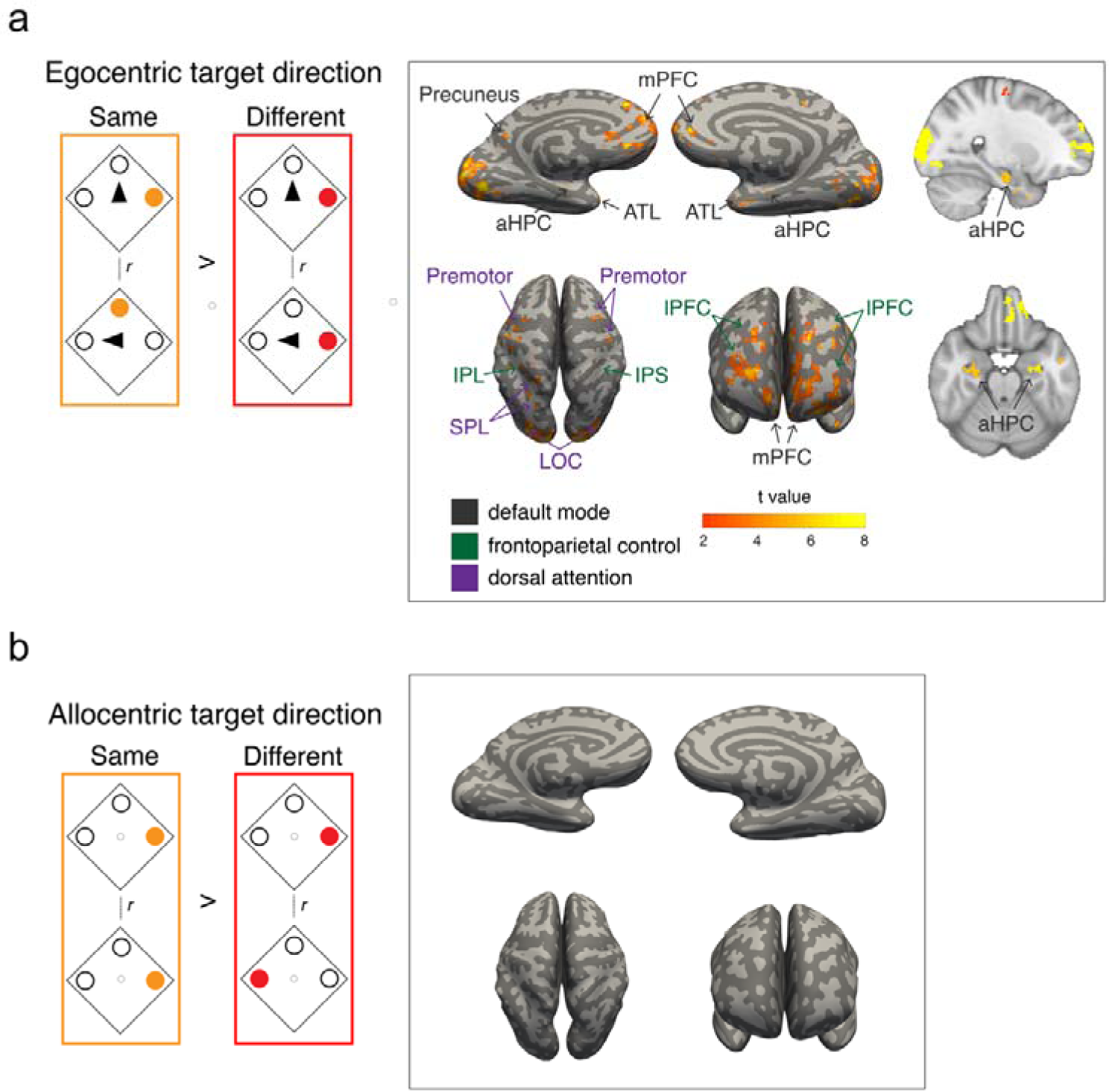
Neural representation of retrieved egocentric target location. (a) Left panel: Schematic representation of decoding the egocentric direction of a target character. Right panel: Clusters were revealed across a wide range of brain areas (P < 0.001, initial threshold; P < 0.05, cluster-corrected for multiple comparison). Many of the clusters belonged to one of the following three functional networks: the default-mode network, frontoparietal control network, and dorsal attention network. The aHPC is shown on sagittal and transverse section of volume image for display purpose (P < 0.001, initial threshold; P < 0.05, cluster-corrected for multiple comparison). (b) Left panel: Schematic representation of decoding allocentric direction of a target character. Right panel: No clusters were revealed even with the use of a liberal threshold (P < 0.01, initial threshold; P < 0.05, cluster-corrected for multiple comparison).

## Increased default-mode network connectivity while locating a target compared with self-locating

The present results showed that the MTL and mPFC signaled a coherent map coding a spatial relationship of the three human characters during the different time periods in which different task demands were required (i.e., self-locating and target-locating). In addition, the MTL and mPFC signaled the different location information even during the same targeting period; MTL areas tended to represent self-orientation while the mPFC tended to represent egocentric target location. To investigate how the different functional contributions of the MTL and mPFC were substantiated by whole brain large-scale networks, we manually segmented the MTL sub-regions in each participant’s native space (Fig. S4a, top panel), and conducted a task-based functional connectivity analysis using each of MTL and mPFC sub-regions as seed (six and four, respectively). For each seed, the mean regional bold signals associated with two TRs in each of facing and targeting period were estimated in each trial, and concatenated across trials to make its task-based time course, which contains 72 time points (i.e., 2 TRs × 36 trials) in each session. The task-based time course of the regional signal for each seed was correlated with each voxel’s time course outside the seed and then was averaged across the four sessions for each participant, generating seed-based connectivity maps for each of facing and targeting periods (Ranganath et al., 2005). Then, we compared the connectivity between the two periods across the participants using a permutation test (see Methods for details).

First, we examined the connectivity between the MTL and mPFC subregions for each of task demands, the result indicated a significantly larger connectivity between the medial orbital frontal cortex and the MTL subregions in the targeting period relative to the facing period. (aHPC: t(18)=4.75, P=0.003, pHPC: t(18)=3.96, P=0.02, PHC: t(18)=6.85, P<0.001, bonferroni corrected for multiple comparison (n=24)). In addition, both the MTL and mPFC showed significantly larger connectivity to brain areas that belong to the default-mode network and those to the dorsal attention network during the targeting period compared to the facing period (Fig. 5b, P < 0.001, voxel-wise threshold; P < 0.05, cluster-corrected for multiple comparison). By contrast, both the MTL and mPFC showed significantly larger connectivity to the frontoparietal control network during the facing period relative to the targeting period (Fig. 5a, P < 0.001, voxel-wise threshold; P < 0.05, cluster-corrected for multiple comparison). These results suggest that both the MTL and mPFC changed their connectivity to the three functional networks depending on the task demands. We next evaluated the task-based functional connectivity during each task period based on the three functional network masks (Fig. S4b). This ROI analysis revealed that the default-mode network was positively correlated with the MTL (t(18) = 7.98 for average across the sub-regions within MTL, P < 0.001) and mPFC (t(18) = 9.63 for average across the sub-regions within mPFC, P < 0.001) for both time periods with a significant increase during the targeting period (Fig. 5c & Fig. S5, top panel; F(1,72) = 4.51, P = 0.03), regardless of the seeds (MTL or mPFC; F(1,72) = 0.00, P=0.98). In contrast, the frontoparietal control network showed significantly negative connectivity with the MTL (t(18) = −10.50, P < 0.001) and mPFC (t(18) = −6.55, P < 0.001) during both task periods, this negative connectivity was stronger during the targeting period (Fig. 5c F(1,72) = 5.58, P = 0.02, also see Fig. S5, middle panel;). These results suggest that the default-mode network contributes more to the retrieval of the target location than the self-location to an external reference. Interestingly, despite both the MTL and mPFC being part of the default-mode network, they showed opposite connectivity patterns to the dorsal attention network during both periods (Fig. 5c & Fig. S5, bottom panel; F(1,72) = 55.07, P < 0.001); the MTL positively with the network while the mPFC negatively correlated with it. The connectivity between the MTL and the dorsal attention network increased from the facing to targeting period (F(1,72) = 8.43, P = 0.005). These results suggested that the dorsal attention network, which contains the superior parietal lobule (SPL) that represented egocentric target location (Fig. 4a), showed increased coupling with the MTL during the targeting period.

**Figure 5:**
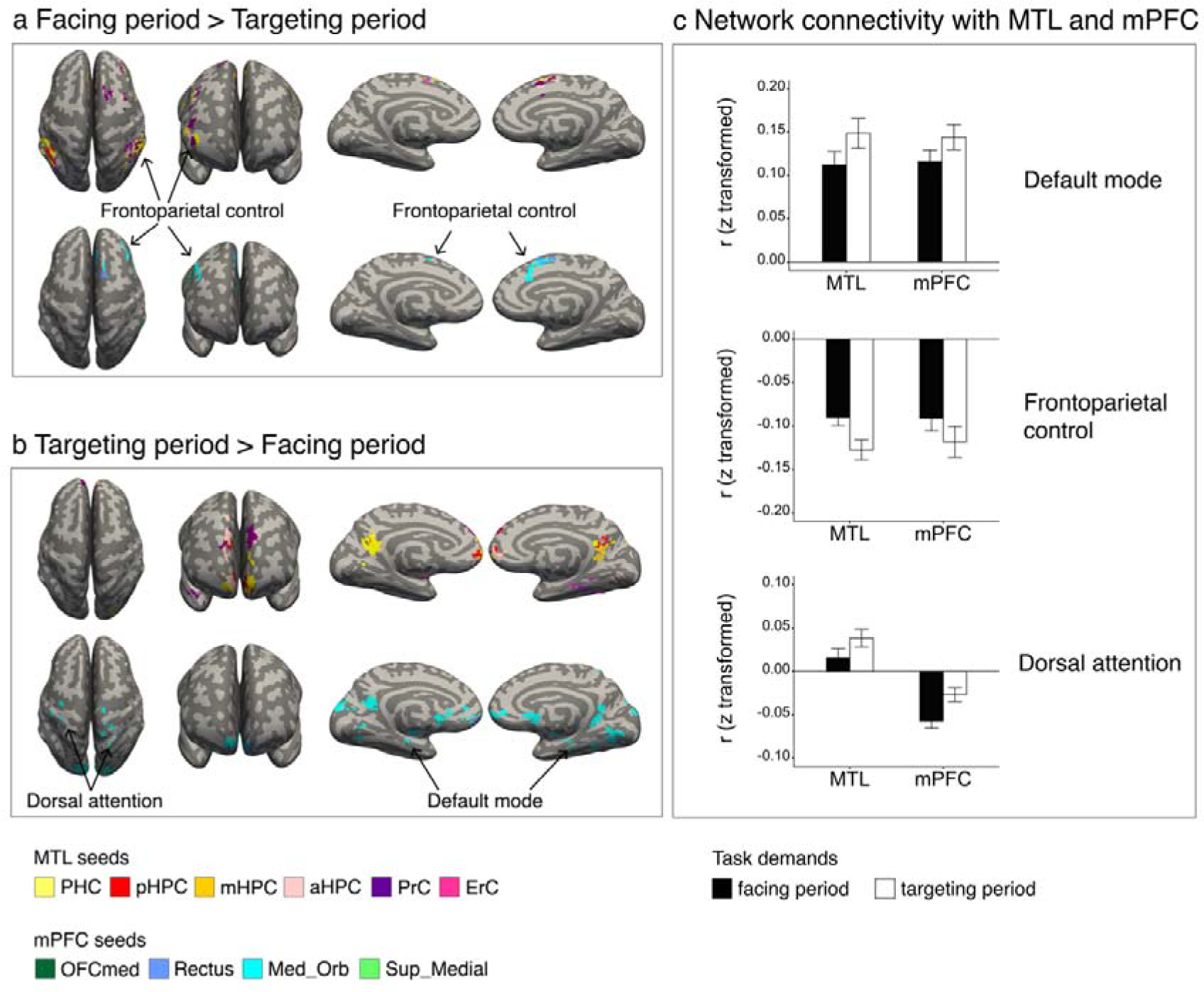
Increased default-mode network connectivity while locating a target compared with locating oneself. (a) The frontoparietal control network showed enhanced connectivity strength with the MTL and mPFC in the facing period compared to the targeting period (P < 0.001, initial threshold; P < 0.05, cluster-corrected for multiple comparison). (b) The default-mode network and dorsal attention network showed enhanced connectivity strength with the MTL and mPFC in the targeting period compared to the facing period (P < 0.001, initial threshold; P < 0.05, cluster-corrected for multiple comparison). (c) The mean connectivity strength of MTL and mPFC sub-regions with three networks, respectively. Note that the connectivity between default-mode network and MTL/mPFC was examined using default-mode network mask without the MTL/mPFC, respectively.

## Discussion

In this study, we examined neural representations of space defined by three objects and found that both the HPC and mPFC represented the object-based space around the participants. Interestingly, the HPC represented the object-based map when the participants locate their self-body in the environment constructed by the three objects, while the mPFC represented the map when the participants remembered the location of a target object relative to the self-body. These results suggest that the cognitive maps in different brain regions play different functional roles. In addition, during the targeting period, we found differential spatial representations across the MTL and mPFC: the MTL generally reinstated self-orientation, while the mPFC represented egocentric target location relative to self-location. Increased functional connectivity was observed between the MTL and mPFC under the necessity of the retrieval of the target location from the stored memory (targeting period) compared to when they actually faced the reference object to locate their self-body (facing period). These results suggest that mental representations of the external world formed by the coherent space and its constituent elements may be shared in the default-mode network including the MTL and mPFC. The special role of the mPFC in this scheme might be to select the object location based on the mnemonic information including the cognitive map and current self-location on it, which might be propagated from the MTL.

To examine the representation of spatial “*maps*” (Fig. 1b), the present task was designed to cancel out effects of a particular encoding experience related with the walking direction as well as a particular object identity that the participants viewed during the facing and targeting periods in each trial by balancing number of trials with each of those confounding factors in each map (see Methods). Therefore, the neural representation of the map information revealed by the RSA could not be explained by perceptual information in the present study. Moreover, the participants always stood on the center of the virtual environment during the facing and targeting periods, during which the map effect was examined. Because of this task design, the map information does not directly indicate self-location information like place fields of place cells in the HPC (O’Keefe and Dostrovsky, 1971). On the other hand, the representations of place-fields are reportedly influenced by the animal’s cognitive map, and the existence of cognitive maps could be most clearly demonstrated by a phenomenon known as “remapping”, which reportedly occurs in populations of place cells in the rodent HPC (Moser et al., 2017). Therefore, it might be reasonable to interpret the map representations in the left mHPC during the facing period as experimental evidence of “remapping” of place cells in the human HPC even though the participants stood in the same position. However, holding this interpretation predicts that human place cells are localized in the left mHPC. This prediction is against consistent evidence from previous human studies reporting that the right HPC was more involved in encoding and retrieving spatial information than the left HPC (Abrahams et al., 1997; Maguire et al., 1997; Ekstrom et al., 2003; Doeller et al., 2008; Schinazi, 2013). The other possible interpretation for the map representation in the left mHPC is that it may encode an allocentric spatial relationship of the three objects itself. This interpretation is consistent with previous human imaging studies reporting contributions of the left HPC to the imagination of visual scenes, which could be constructed from multiple spatial elements (Addis et al., 2007; Bird et al., 2010). The specific role of the left HPC in relational memory was also reported in non-spatial information domains, including associative learning (Kumaran et al., 2009; Suarez-Jimenez et al., 2018) and social interactions (Tavares et al., 2015;). Taken together, it might be more reasonable to interpret that the clusters in the left mHPC was related to a coherent space constructed by the multiple objects rather than its influence on representations of individual spatial elements such as self-location or head direction. RSA also suggested the involvement of the PRC and PHC in MTL signaling the object identity and egocentric view of their spatial layout, respectively, which might be used for constructing the coherent map from its constituents in the left mHPC. Future studies should address how the coherent map can be constructed by multiple objects in the MTL.

In contrast to the facing period, the representation of the map information was found in the mPFC but not in the HPC during the targeting period. In addition, the mPFC signaled the egocentric location of a targeting object, while the MTL concurrently signaled the self-orientation. Involvement of the mPFC in constructing goal-directed information in the current context is consistent with accumulating evidence showing that the mPFC contributes to decision making or action selection (Saxena et al., 1998; Gallagher et al., 1999; Feierstein, 2006; Spiers and Maguire, 2007; Kable and Glimcher, 2009; Young and Shapiro, 2011; Balaguer et al., 2016; Yamada et al., 2018). These previous studies consistently supported the notion that the mPFC function becomes obvious when an appropriate selection requires mnemonic information in addition to incoming perceptual information (Bradfield, 2015). In this study, together with perceptual information responsible for target object identity, mnemonic information such as the map information and self-orientation was required to solve the task. Considering that the MTL could provide all the necessary mnemonic information, a reasonable interpretation is that the mPFC was involved in the selection of a target location among alternatives rather than the recollection or generation of it.

In addition to the HPC and mPFC, the map information has been observed in other brain areas such as the angular gyrus (Seghier, 2013; Price et al., 2016), lateral temporal gyrus (Karnath, 2001; Himmelbach et al., 2006), and precuneus (Cavanna and Trimble, 2006) that also belong to the default-mode network. The brain areas in the default-mode network, particularly the MTL sub-regions except for the PRC represented self-orientation during the targeting period. On the other hand, RSA analysis showed that the representation of the egocentric target object location recruited widely-distributed brain regions, which belong not only to the default-mode network but also to the dorsal attention network and frontoparietal control network. The increased functional connectivity between the MTL and the brain regions of dorsal attention network as well as the mPFC suggest that the egocentric target location signal is transmitted from the mPFC to the dorsal attention network, such as the SPL (Evans et al., 2016), via the MTL, which implies a pivotal functional role of MTL as a hub of mental representation of object-related (e.g., identity and location) signals. This signal transmit across different brain networks may be related with the fact that the egocentric target location was the main behaviorally relevant feature in the present task. These results contrast with the present result that no brain regions represented the allocentric target location relative to the spatial layout of the characters, which was not required to answer in the task.

Interestingly, the frontoparietal control network showed a strong negative correlation with both the MTL and mPFC during both the facing and targeting periods, although the lPFC in the frontoparietal control network represented both the map information and egocentric direction during the targeting period. In addition, the lPFC represented walking direction as well as character identity during both periods. These results suggest that the lPFC computes the target location independently of the default-mode network. The parallel contributions of the lPFC and MTL-mPFC in choosing the target location may reflect their different cognitive functions (Jimura et al., 2004). lPFC has long been considered as a center of executive functions (Funahashi, 2017; Miller et al., 2018) equipped with working memory (Andrews et al., 2011; Barbey et al., 2013; Brunoni and Vanderhasselt, 2014; Funahashi, 2017). In human fMRI studies, the lPFC has been shown to contribute to the retrieval of task-relevant information when more systematic thinking is required (Epstein et al., 2017; Javadi et al., 2017). In the present study, the behavioral task was designed to ensure participants neither actively maintained a spatial configuration of the human characters during the walking period nor any systematic strategy to solve the task, which was confirmed by the post-scanning test results. The greater signal for the cognitive map and the egocentric target location in the mPFC than that in the lPFC may reflect that the current spatial memory task was enough easy to allow participants to depend only on the involuntary encoding and subsequent memory retrieval for their top-ceiling performance (Epstein et al., 2017; Javadi et al., 2017).

In contrast to previous memory/navigation studies, which examined brain functions using spatial environments consisting of immobile landmarks (e.g., stores) and/or landscapes (e.g., mountains) (Bird et al., 2010; Woollett and Maguire, 2011; Schinazi et al., 2013; Chadwick et al., 2015; Brown et al., 2016), the present study used a spatial environment constructed by only mobile objects that could become targets and references of self-location as well as determine the space (i.e., map) around oneself. This task design allowed us to extract a mental representation of the spatial environment consisting of the minimum essential constituents. This reductionist method could be useful for future studies investigating the construction and functional mechanisms of a cognitive map because of its simplicity. One critical concern might be whether the findings discovered by this reductionist method can be applied to a more complicated cognitive map consisting of large numbers of immobile spatial elements, which could be learned through extensive explorations over a long time period (e.g., the city of London) (Woollett and Maguire, 2011). Another related concern might be whether our brain system holds only one cognitive map or multiple ones at a time (Meister and Buffalo, 2018). For example, we may hold an object-based cognitive map consisting of relevant mobile objects such as same species, predators, and foods, while we may also hold the other cognitive map consisting of landmarks, landscapes and other immobile objects such as trees. Future studies should address the relationships of different types of cognitive maps (e.g., mobile vs immobile, short-term vs. long-term) and their underlying neural mechanisms.

The present study found neural representations of the space specified by objects around us. This object-based cognitive map seems to interact with representation of self-location in HPC and to mediate a selection of egocentric target-location in mPFC, which would serve for leading us to the goal position. In addition to the spatial navigation, an existence of the object-based cognitive map may equip us with a space representation for persons separately from the background, which may serve for our social interactions (Damasio et al., 1994; Stolk et al., 2015) as well as the encoding and retrieval of episodic memory (Tulving, 2002; Squire and Wixted, 2011).

## Supplemental Information

Supplemental Information includes six figures, two tables and two videos. The videos contain trial examples for Day 1 and Day 2. In the video of day 1, please note we only used examples of correct trials in which a green square of line was preented as feedback after the participant’s response. File size: 21.5 MB; Video duration: 1.23 minute; File format: .mp4; Video codec: H.264; Aspect ratio: 1024 × 768.

## Author Contributions

B.Z. and Y.N made the experimental design; B.Z. conducted all experiments and data analysis under supervision of Y.N.; B.Z. and Y.N. wrote the paper.

## Acknowledgments

The present study was funded by National Natural Science Foundation of China Grant 31421003 (to Y.N.). Computational work was supported by resources provided by the High-performance Computing Platform of Peking University. We thank Li Sheng and Sun Pei for helpful discussions, we also thank Arielle Tambini, Lusha Zhu, Koji Jimura, Rei Akaishi and Cen Yang for comments on an early version of the manuscript.

## Materials and Methods

### Participants

Nineteen right-handed university students with normal or corrected-to-normal vision were recruited from Peking University (12 females, 7 males). The average age of the participants was 24.9 years (range: 18–30 years). All participants had no history of psychiatric or neurological disorders and gave their written informed consent prior to the start of the experiment, which was approved by the Research Ethics Committee of Peking University.

### Experimental design

#### Virtual environment

We programmed a 3D virtual environment using Unity software (Unity Technologies, San Francisco). The environment was designed with a circular fence as a boundary (48 virtual meters in diameter), a flat grassy ground, a uniform blue sky, and with a wood plate surrounded by four vertices of a square placed in the center (Fig. 1b, 4.7 virtual meters for side length). Three human characters (Mixamo, San Francisco, https://www.mixamo.com) were placed on three of the vertices in each trial. A map was defined by the relative relationship of the three human characters (Fig. 1b). From the six possible maps, three of them were pseudo-randomly selected for each participant to collect enough number of trials’ data for each condition during the allowable range of scanning duration. The maps were the only environmental cues relevant to the task requirement, no distal cues were used outside the boundary. Participants performed the task using the first-person perspective with a 90° field of view (aspect ratio = 4:3), they had never seen a top-down view of the virtual environment.

#### Walking period

Participants walked from one of four starting locations near the circular boundary (4 virtual meters from the boundary) toward the human characters (Fig. 1c) and stopped on the wood plate. The visual stimuli (spatial environment viewed from first-person perspective) were determined by the combination of the map and walking direction, in other words, each map was presented by four different visual stimuli that were determined by the starting position (Fig. 1c). Importantly, participants were blinded to the map concept throughout the task. The walking period lasted for 6.0 s, during which each character had a 20.6% probability of nodding its head at a random time point between the start and end of walking. There was a 50%, 38.9%, 10.2%, and 0.9% probability for 0, 1, 2, and 3 characters to nod head in each trial; we subjectively selected a 20.6% head-nodding probability for each character to ensure an approximately equal number of trials with head-nodding and no head-nodding. During the walking period, participants were required to pay attention to the heads of the human characters rather than to memorize their spatial arrangement. The height of the participants was 1.8 virtual meters from the ground, which was the same as that of the human characters. No response was required during the walking period.

Two tasks were completed in two consecutive days. On day 1, the participants performed an HND task that did not include spatial-memory trials. On day 2, participants performed a spatial-memory task.

#### Head-nodding detection (HND) task

Participants performed 144 randomly ordered HND trials in a behavioral experimental room. In each trial, a photo of one of the characters was presented on a screen after the walking period, and participants were asked to indicate whether the character nodded its head or not (Fig. S1a). For this task, there was a 50% chance that the character in the presented photo nodded its head. Feedback was given after the participants had responded with either green (correct) or red (incorrect) photo border. The stimuli were rendered on a PC and presented on a 27-inch LCD monitor (ViewSonic XG2730) with a screen resolution of 1024 × 768. The HND task was used to examine whether participants paid attention to head-nodding rather than memorizing the spatial arrangement of the characters, which would be indicated by high success rates in the head-nodding test.

#### Spatial-memory task

During this task, participants performed 144 spatial-memory trials (90%) and 16 HND trials (10%) that lasted ∼ 70 min in an MRI scanner. Participants were notified that the remuneration depended only on the performance in the HND trials although they were also encouraged to perform the spatial memory task as best as they could (videos of trial examples are available online for both tasks). The trial-type (i.e., HND or memory task) was distinguishable after the walking period by subsequent stimuli. In the spatial-memory task, participants experienced a “facing period” and a “targeting period” sequentially after the walking period. In the facing period, their self-orientation was changed immediately to one of the human characters (facing-character) without viewpoint transition, and a character with the environment background was presented for 2.0 s with the other two characters being invisible, the participants were instructed to face the character. In the targeting period, a photo of another character (targeting-character) was presented as a target on a scrabbled background for 2.0 s. Each of the three experimental periods was followed by a 2.0-s delay (noise screen). At the end of each trial, participants indicated the direction of the target character relative to their self-body by pressing a button when a cue presented on the screen; no feedback was shown for both trial-types (Fig. 1a). The spatial-memory task contained four experimental sessions, each containing a spatial information combination of 3 maps × 4 walking directions × 3 facing-character identities in each session, with targeting-characters balanced across sessions. After scanning, all participants completed a post-scanning interview and reported the strategy they used to perform the task (Table. S2).

### fMRI data acquisition

Imaging data were collected using a 3T Siemens Prisma scanner equipped with a 20-channel receiver head coil. Functional data were acquired with a Multi-band Echo Planer imaging (EPI) sequence (TR = 2000 ms, TE = 30 ms, matrix size: 112 × 112 × 62, flip angle: 90°, gap = 0 mm; resolution: 2 × 2 × 2.3 mm^3^, number of slices: 62, slice thickness: 2 mm, slice orientation: transversal), four experimental sessions were collected with, on average, 478, 476, 473, 475 TRs, respectively. A high-resolution T1-weighted three-dimensional anatomical data set was collected to aid in registration (MPRAGE, TR = 2530 ms, TE = 2.98 ms, matrix size: 448 × 512 × 192, flip angle: 7°, resolution: 0.5 × 0.5 × 1 mm^3^, number of slices: 192, slice thickness: 1 mm, slice orientation: sagittal). During scanning, experimental stimuli were presented through a Sinorad LCD projector (Shenzhen Sinorad Medical Electronics) onto a 33-inch rear-projection screen located over the subject’s head with a resolution of 1024 × 768 and viewed with an angled mirror positioning on the head coil.

### fMRI preprocessing

Functional data for each session were preprocessed independently using FSL FEAT (FMRIB’s Software Library, version 6.00, https://fsl.fmrib.ox.ac.uk/fsl/fslwiki; Woolrich et al., 2001; Woolrich et al., 2004). For each session, the first three functional volumes were discarded to allow for T1 equilibration, and the remaining functional volumes were slice-time corrected, realigned to the first image, and high-pass filtered at 100 s. For group-level statistics, each session’s functional data were registered to a T1-weighted standard image (MNI152) using FSL FLIRT (Jenkinson and Smith, 2001), and this procedure also resampled the functional voxels into a 2 × 2 × 2 mm resolution. For RSA, data were left unsmoothed to preserve any fine-grained spatial information (Chadwick et al., 2012). For functional connectivity analysis, data were smoothed using a 5 mm FWHM Gaussian kernel and were high-pass filtered at 0.01 Hz to remove low-frequency signal drifts.

### Anatomical masks

We manually delineated the MTL, including the HPC, PHC, PRC, and ERC on each participant’s native space using established protocols (Insausti et al., 1998; Pruessner et al., 2000; Pruessner et al., 2002; Duvernoy, 2005), as well as a delineating software ITK-SNAP (www.itksnap.org). The HPC was further divided into its anterior, middle, and posterior parts given the anatomical and functional variability along the HPC long axis (Poppenk et al., 2013), the anterior border of pHPC and the posterior border of aHPC were defined by the appearance of the crus of the fornix and the uncal apex relative to mHPC along the coronal orientation, respectively (Pruessner et al., 2000; Poppenk et al., 2013). For PFC sub-regions, we used the AAL template (Rolls et al., 2015), and selected four mPFC sub-regions for ROI-analysis, which included the rectus, medial orbital gyrus (OFCmed), medial orbital frontal gyrus (Med_Orb), and superior medial frontal gyrus (Sup_Med). All ROIs were resampled and aligned with the functional volumes, and voxels outside of the brain were excluded.

### Representational similarity analysis (RSA)

Task-relevant information was decoded using RSA. We tried to dissociate the neural effect of facing and targeting period based on 4 s duration from each period onset to the end of following noise period (Zeithamova et al., 2017). First, the trial-based multi-voxel activity patterns of two periods were obtained by creating two separate univariate general linear models (GLM). In each GLM, the 4 s blood-oxygen-level-dependent (BOLD) signals of 36 trials (a session) were modeled using boxcar regressors. In addition to the 36 trial-based regressors of interest, nuisance regressors were included, which included twelve regressors for modeling the visual patterns of the walking period determined by the maps and walking directions, three for modeling the character identities in the remaining period (for example, in the facing period GLM, three targeting characters were specified as nuisance regressors), four for modeling head-nodding detection trials, three for modeling 3 directional cues in the response period, and six motion parameters. This procedure generated 36 trial-based multi-voxel patterns in participant’s native space (2 × 2 × 2.3 mm voxels) for each period, those multi-voxel patterns were normalized prior to subsequent analysis by subtracting the grand mean pattern of the 36 multi-voxel patterns for each session (Vass and Epstein, 2013).

Searchlight-based RSA. Next, we computed the representational similarity for each spatial information based on the 36 multi-voxel patterns using a searchlight-based RSA (Libby et al., 2014; Chadwick et al., 2015), which was conducted using custom Matlab (version R2018b, www.mathworks.com/matlab/) scripts. In detail, a sphere with a 6 mm radius was constructed (85 voxels per sphere) for each brain voxel, and the spheres near the edge of the brain with fewer than ten voxels were excluded from the analysis. The activity parameters within each sphere were extracted from each of the 36 multi-voxel patterns, resulting in a 36-column by n-row (number of voxels within the sphere) matrix. The pattern similarity was then calculated between each column-by-column pair using Pearson’s correlation, and was normalized using Fisher’s r-to-z transformation. This procedure finally generated a 36-by-36 correlation matrix for each period in each brain voxel. Next, given that a multi-voxel pattern contains the combination of multiple spatial information, we conducted a GLM as a regression on the correlation matrix, which included each element from one side of diagonal of the matrix but did not the diagonal elements, by specifying multiple categorical regressors to dissociate an effect of each spatial information form potential influences of others. In each regressor, either “1 (same)” or “0 (different)” were used to correspond with the correlation coefficient of a given column-to-row element of the correlation matrix. An estimated parameter of each regressor evaluated an increase or decrease in the similarity in the same condition relative to the different condition concerning on the spatial information specified by the regressor. For the facing period, the GLM contained five categorical regressors, which included the (1) “map”, (2) “walking direction”, and (3) “facing-character identity”. Since the participants reported thinking about their bodies rotating between the walking direction and self-orientation relative to the environment, we also added the (4) “rotation angle” (turn left/right 45°, turn left/right 135°), and (5) their “self-orientation” into the GLM. For the targeting period, seven regressors were built, which included: (1) “map”, (2) “walking direction”, (3-4) participants’ “rotation angle” and “self-orientation”, (5) “targeting-character identity”, and (6-7) “egocentric and allocentric position of target-character”. It is important to note that the “facing-character identity” was not included in the targeting period GLM since the effect of each facing character was regressed out in the GLM computing of multi-voxel activity patterns. r^2^ was computed and ranged from 0 to 0.03 for the facing period GLM, and 0 to 0.04 for the targeting period GLM (Table S1). Each regressor’s parameter was then assigned to the center voxel of each sphere so that a whole-brain statistical parametric map could be generated for each spatial information for each period, with those spatial representations being finally averaged across the four scanner sessions.

#### ROI-based RSA

To validate the representation of the cognitive map information in the HPC and mPFC suggested by the searchlight-based RSA, we further conducted an independent RSA using anatomical ROIs for the both hemispheres of mHPC and rectus. We reasoned that since searchlight analysis identifies the spatial representations as clusters in small portions of anatomical regions, if those representations are stable enough, the corresponding anatomical regions, on average, should show a clear increasing tendency in similarity when the to-be-tested spatial information are same in trial-pairs compared to a chance level such that they match the searchlight results. To test this, we separated each of mHPC and rectus into the left and right hemispheres and generated 4 anatomical ROIs, the rectus masks were normalized into the participants’ native space. For each ROI, the multi-voxel pattern similarity was calculated independently for “same-map” and “different-map” conditions from the correlation matrix of each period, then the similarity of both conditions was averaged across the four sessions for each participant. The chance level was determined by permutation test, in which the trial labels on the correlation matrix were shuffled before calculating the representational similarity of “same-map” and “different-map” conditions. This procedure was performed by 5000 times for each ROI, the chance level was obtained for each condition by averaging the 5000 shuffled similarity results. We then subtract the chance level from the similarity in each condition, and tested whether or not the baseline-corrected similarity is significantly different from zero across the participants using one-sample t-test (two-tailed).

### Functional connectivity (FC) analysis

To investigate the functional networks for different task demands, we examined the whole-brain FC using each sub-region of MTL and mPFC as seed (Fig. S4a; 12 for the MTL and 8 for the mPFC). In detail, we first removed the nuisance covariates from the preprocessed functional data by creating a GLM, which specified the signal averaged over the lateral ventricles, white matter, and whole brain, six motion parameters, and their derivatives as regressors. The residual signal was bandpass-filtered, leaving signals within the frequency range 0.01 to 0.1 Hz, and was shifted by two TR intervals (4 s) for subsequent analysis (Tompary and Davachi, 2017). We computed a regional time course for each anatomical mask in each of facing and targeting period. To do this, we averaged signals over the mask at each TR within the period, and then concatenated the two TRs in one trial with those in the next trial within a session (Ranganath et al., 2005). The regional time course for each anatomical mask was correlated with the time course of each voxel in the rest of the brain, resulting in a whole-brain correlation map for each period in each scanning session. The correlation maps were averaged across four scanning sessions for each participant, and were then submitted to a two-tailed t-test for group level statistics.

Each cluster, which derived from the contrast analysis in connectivity between facing and targeting period based on an initial threshold of p=0.001, was assigned to each of the three networks based on previous literatures: default-mode network, frontoparietal control network, and dorsal attention network (Vincent et al., 2008; Schacter et al., 2012; Gelström and Graziano., 2017). In the present study, default-mode network contains the clusters of mPFC, MTL, posterior cingulate cortex, and anterior temporal gyrus; frontoparietal control network contains the clusters of paracingulate gyrus, lateral PFC, and inferior parietal lobule; dorsal attention network contains the clusters of occipital pole, lateral occipital cortex, cuneal cortex, lingual gyrus, superior parietal lobule, and postcentral gyrus (Fig. S4b). To examine modulation effects on the connectivity of MTL/mPFC with the large-scale networks by different task demands, we computed mean connectivity between each of MTL and mPFC subregions with each network. For the default-mode network, we prepared for two masks, in which the MTL or mPFC was removed for the examinations of its connectivity with the default-mode network.

### Statistical analysis

For searchlight-based RSA, we used an initial threshold of p < 0.001. If no clusters were revealed, a liberal threshold of p < 0.01 was used. For whole brain FC analysis, an initial threshold of p<0.001 was used to identify robust network patterns. The reliability of clusters was tested using a non-parametric statistical inference that does not make assumptions about the distribution of the data (Nichols and Holmes, 2002; Winkler et al., 2014; Chadwick et al., 2015), the test was conducted with the FSL randomise package (version v2.9, http://fsl.fmrib.ox.ac.uk/fsl/fslwiki/Randomise), and performed 5000 random sign-flips on whole brain searchlight beta images or FC images, we then reported clusters with the size higher than 95% of the maximal suprathreshold clusters in permutation distribution. ROI-based RSA used two-tailed one sample t-test to examine the significance of each anatomical mask. Paired t-test was used to examine the difference in connectivity among MTL and mPFC subregions between task demands. All t-test results were Bonferroni-corrected for multiple comparison. The statistical significance was determined according to whether the corrected *P* value was smaller than 0.05. Analysis of variance (ANOVA) was used to test the influence in connectivity between MTL/mPFC and large-scale functional network along time periods.

---

**Figure S1:**
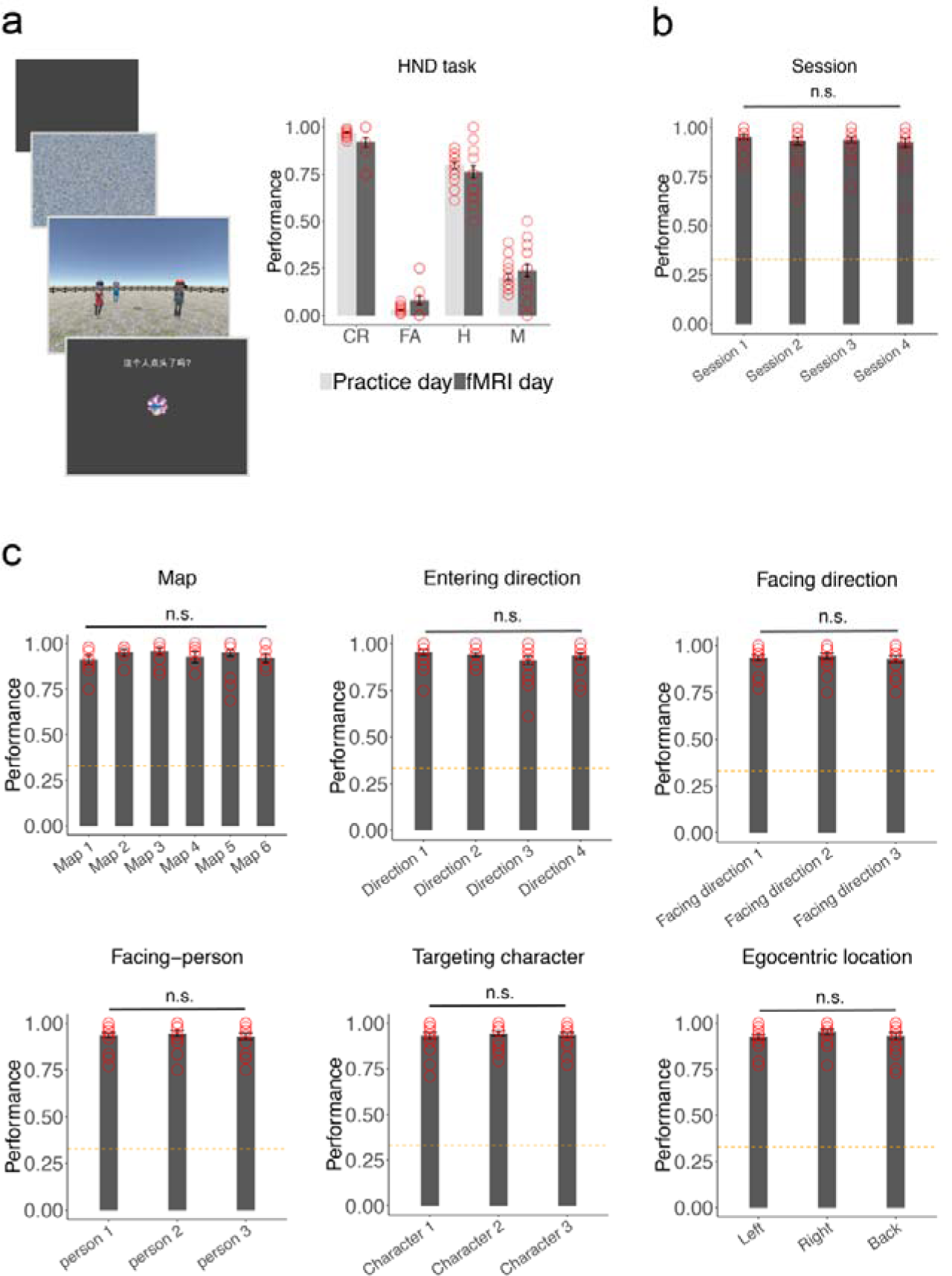
Behavioral results. (a) Paradigm and performance for the head-nodding detection (HND) task. In both the practice and fMRI day, participants exhibited a high accuracy of detecting head nodding and rejecting if there was no head nodding (practice day: head-nodding trials, t(18) = 15.714, P = 5.907 × 10^−12^, no head-nodding trials, t(18) = 112.4, P = 2.2 × 10^−16^; fMRI day: head-nodding trials, t(15) = 7.399, P = 2.224 × 10^−6^, no head-nodding trials, t(14) = 6.315, P = 1.908 × 10^−5^). (b) Performance in the spatial-memory task. No significant difference was found among four sessions (F(3,72) = 0.38, P < 0.001). (c) Performance in the spatial-memory task for each category of spatial information. No significant difference was found among the spatial information in each category (map: F(5,33) = 2.19, P = 0.08; walking direction: F(3,72) = 1.10, P = 0.35; facing character: F(2,54) = 0.27, P = 0.76; facing direction: F(2,54) = 0.94, P = 0.40; targeting character: F(2,54) = 0.15, P = 0.86; egocentric direction: F(2,54) = 0.82, P =0 .45).

**Figure S2:**
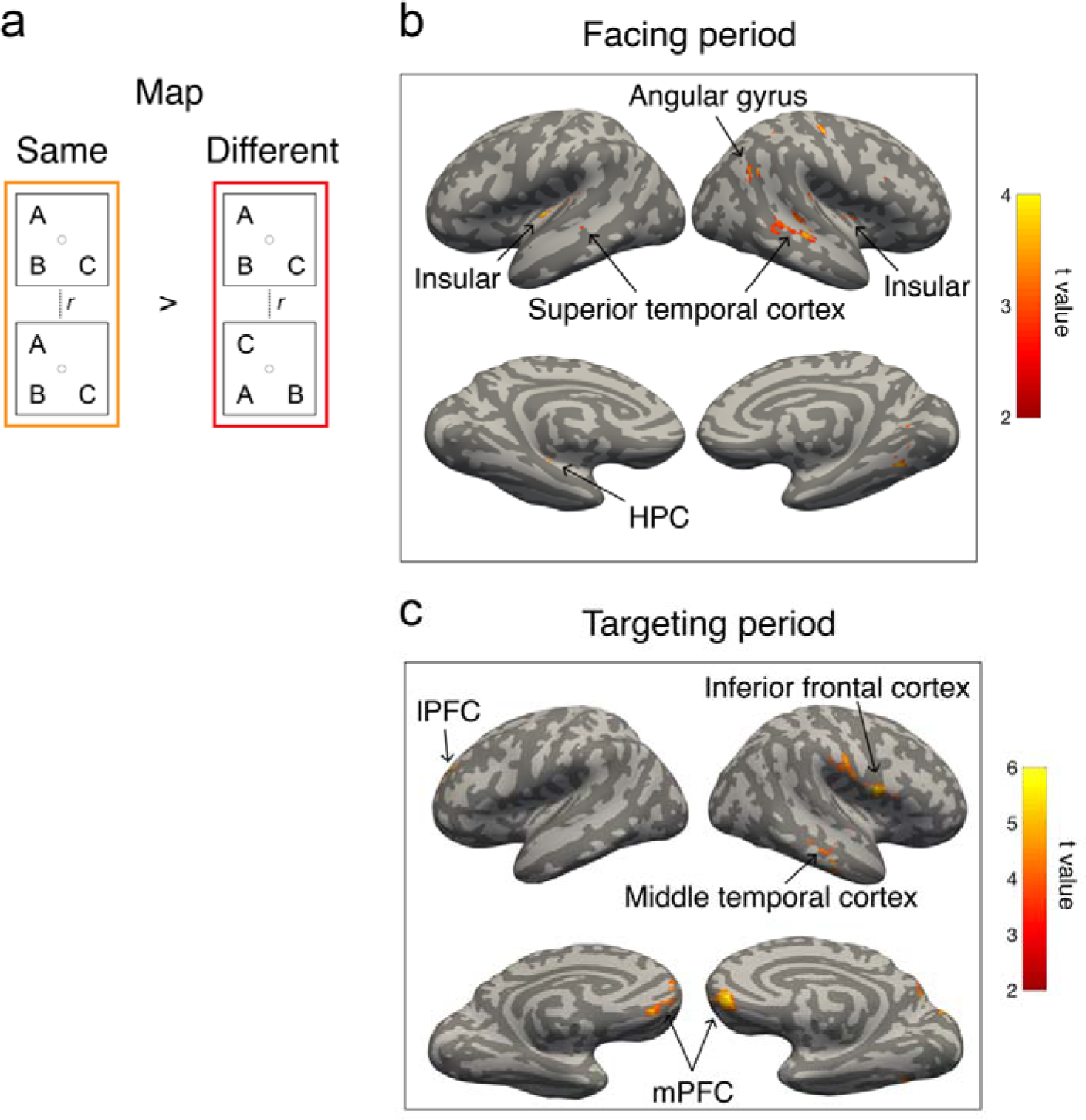
Neural representations of the cognitive map outside of MTL and mPFC, related to Figure 2. (a) Schematic representation of decoding the cognitive map using RSA. (b, c) Neural representation of the cognitive map during facing period (P < 0.01, initial threshold; P < 0.05, cluster-corrected for multiple comparison) and targeting period (P < 0.001, initial threshold; P < 0.05, cluster-corrected for multiple comparison), respectively.

**Figure S3:**
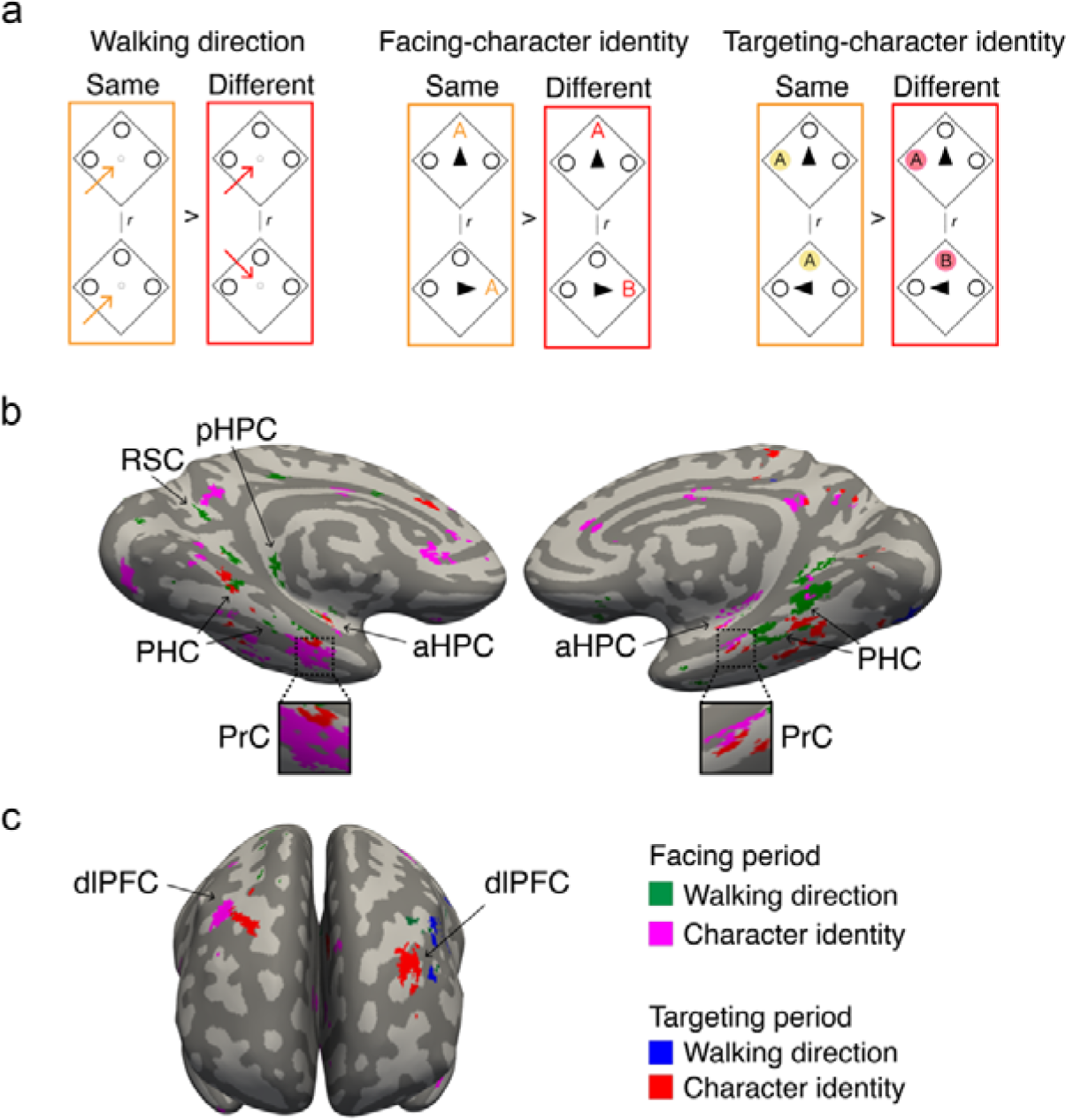
Neural representation of the walking direction and character identity in the MTL, related to Figure 2. (a) Schematic representation for decoding walking direction (left), the character identity in the facing period (middle) and targeting period (right), respectively. (b) The PrC selectively encodes the character identity not the walking direction, and the PHC, PPA, and HPC encodes both the character identity and walking direction. In particular, the PrC encoded both the facing- and targeting-character across both periods, while there was a clear attenuation in the encoding for the walking direction in the targeting period compared to the facing period (P < 0.001, initial threshold; P < 0.05, cluster-corrected for multiple comparison). (c) The clusters located in the lPFC for both the walking direction and character identity across the two temporal periods.

**Figure S4:**
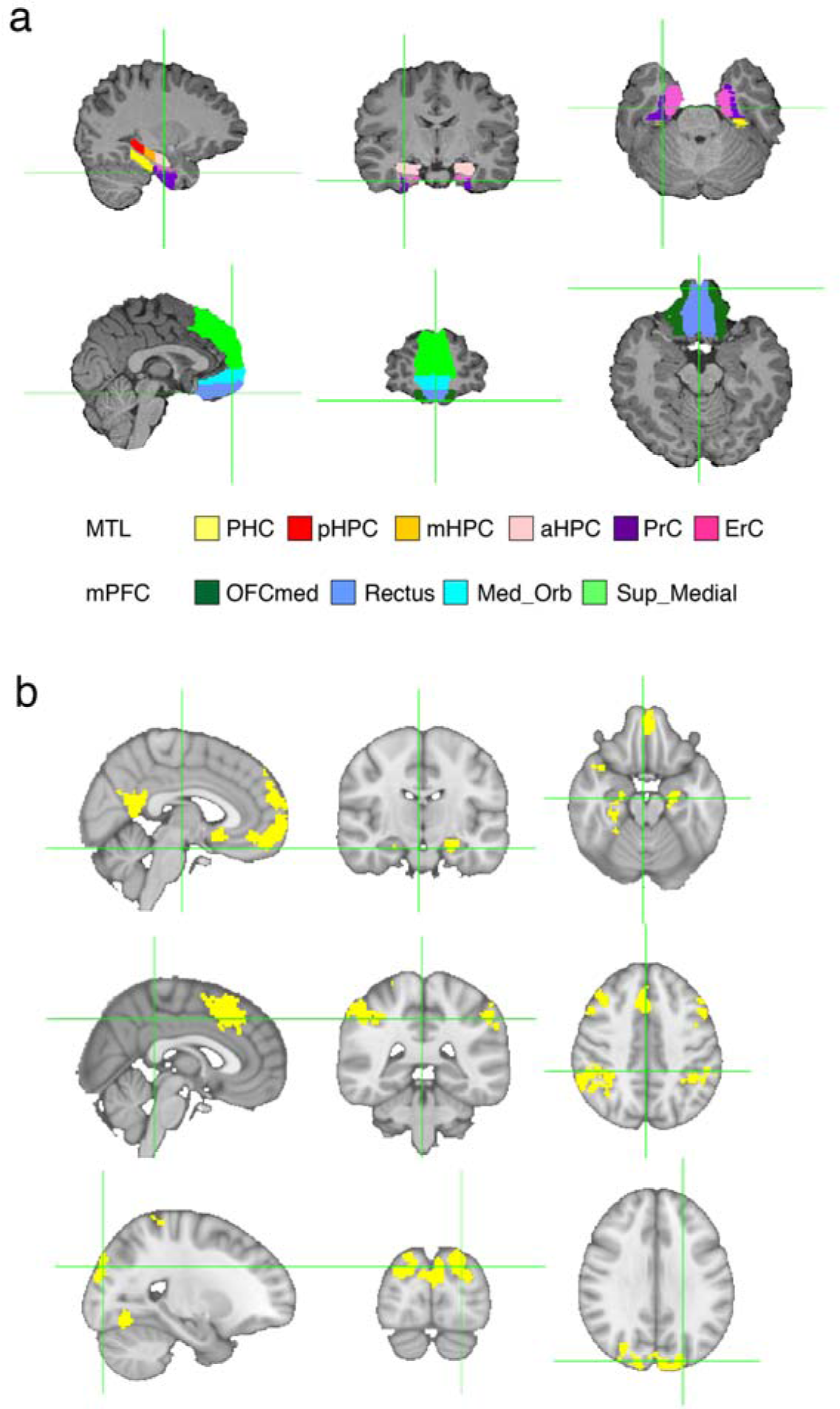
Example of anatomical and functional masks of a participant (No. 13), related to Figure 3, Figure 5 and Figure S5. (a) medial temporal lobe sub-regions (top panel) and mPFC sub-regions (bottom panel) in native space of participants, (b) The network masks identified from the contrast analysis in connectivity between facing and targeting period and were overlapped on MNI152 template. Default-mode network (top panel), Frontoparietal control network (middle panel), Dorsal attention network (bottom panel).

**Figure S5:**
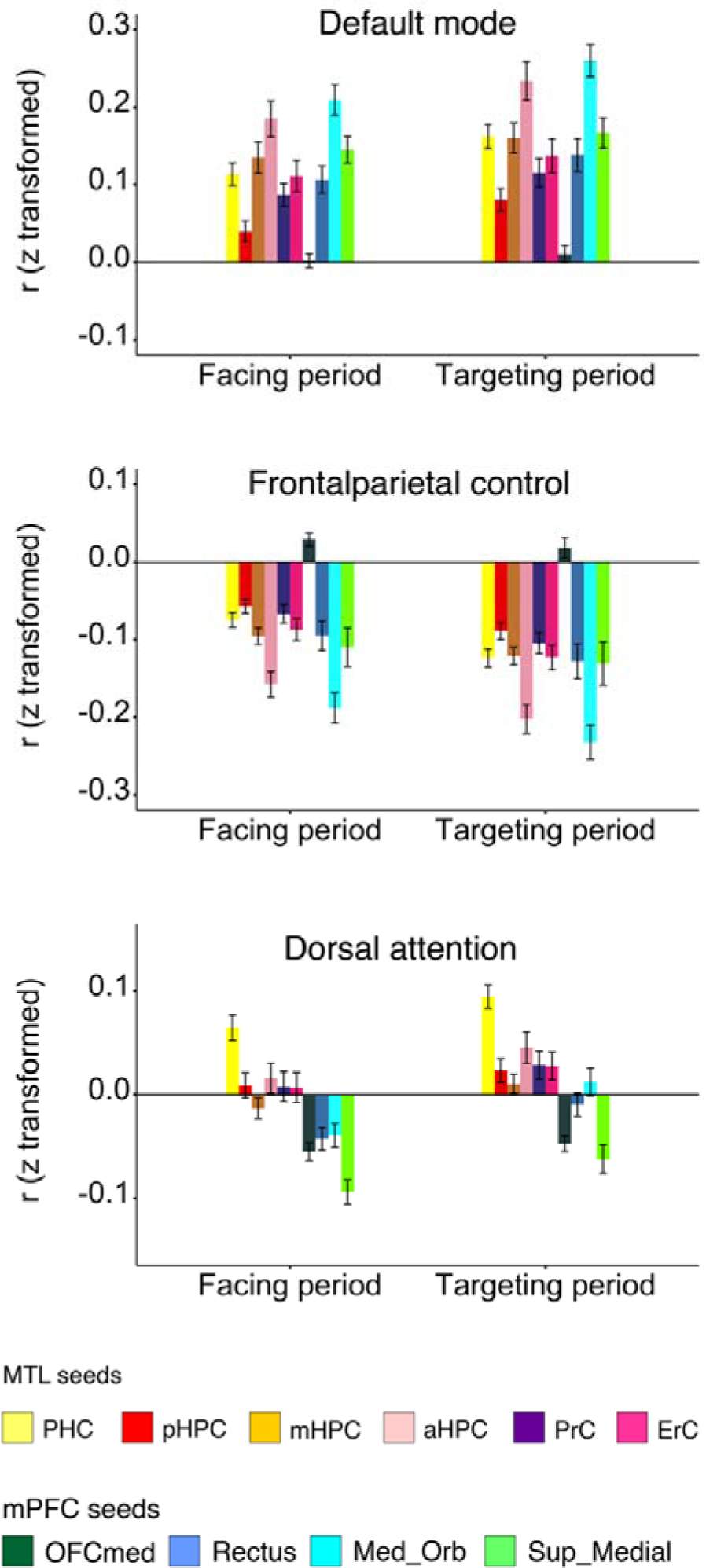
Connectivity strength of the default-mode network, frontoparietal control network, and dorsal attention network with each MTL and mPFC sub-region, related to Figure 5. Note that for the connectivity between default-mode network and MTL/mPFC, two network masks were prepared with the MTL and mPFC excluded, respectively.

**Table S1:** Averaged r2 among searchlight-based GLM regressors for the facing period (top) and the targeting period (bottom) across participants, related to Figure 2, Figure 3, and Figure 4.

|  | Map | Walking direction | Facing character identity | Rotation angle | Self-orientation |
| --- | --- | --- | --- | --- | --- |
| Map | 1.0000 | 0.0057 | 0.0037 | 0.0057 | 0.0037 |
| Walking direction | 0.0057 | 1.0000 | 0.0057 | 0.0008 | 0.0057 |
| Facing character identity | 0.0037 | 0.0057 | 1.0000 | 0.0035 | 0.0307 |
| Rotation angle | 0.0057 | 0.0008 | 0.0035 | 1.0000 | 0.0035 |
| Self-orientation | 0.0037 | 0.0057 | 0.0307 | 0.0035 | 1.0000 |

|  | Map | Walking direction | Targeting character identity | Self-orientation | Allocentric position of target | Egocentric position of target | angle |
| --- | --- | --- | --- | --- | --- | --- | --- |
| Map | 1.0000 | 0.0057 | 0.0037 | 0.0037 | 0.0037 | 0.0057 | 0.0037 |
| Walking direction | 0.0057 | 1.0000 | 0.0000 | 0.0057 | 0.0000 | 0.0008 | 0.0000 |
| Targeting character identity | 0.0037 | 0.0000 | 1.0000 | 0.0000 | 0.0307 | 0.0000 | 0.0000 |
| Self-orientation | 0.0037 | 0.0057 | 0.0000 | 1.0000 | 0.0418 | 0.0035 | 0.0418 |
| Allocentric position of target | 0.0037 | 0.0000 | 0.0307 | 0.0418 | 1.0000 | 0.0000 | 0.0418 |
| Egocentric position of target | 0.0037 | 0.0000 | 0.0000 | 0.0418 | 0.0418 | 0.0000 | 1.0000 |
| angle | 0.0057 | 0.0008 | 0.0000 | 0.0035 | 0.0000 | 1.0000 | 0.0000 |

**Table S2.** Post-scanning interview questions (1^st^ column), Typical feedback summarized from participants (2^nd^ column), Percentage of typical feedback for each question relative to all participants (3^rd^ column).

| Question | Typical feedback | Percentage |
| --- | --- | --- |
| What did you direct your attention to during walking? | I focused on the heads of 3 characters to detect the head-nodding. | 100% |
| Did you try to memorize the relative position of 3 characters?<br>If yes, how did you memorize?<br>If no, how did you solve the task? | No, I solved the task because I remembered the scene that I had looked at during the walking period. | 79% |
|  | Yes, I tried to memorize each scene like photograph during the walking period. | 21% |
| Do you think whether you walked toward 3 characters in the same direction across all trials or in different directions? | I moved in same direction for all the time. | 89% |
|  | It might be different directions although I am not sure. | 11% |
| Did you notice how many different spatial arrangements of 3 characters were presented during the task? If yes, answer the number of spatial arrangements. | No, I did not notice the number of arrangements. | 100% |
| Did you feel as your body rotated or as a spatial environment rotated around you when you looked at the facing character? Do you think whether or not the angles differed across trials? | It was my body rotating, I felt the difference in rotation angle across trials. | 84% |
|  | It was my body rotating, I did not notice the difference in the rotation angle. | 16% |
| What did you think about when you looked at the facing-character? | I recalled the relative position of the three characters during walking, and my position and orientation after walking. | 100% |
|  | I imagined the three human characters' directions relative to me. | 79% |
| What did you think about when you looked at the targeting character? | I first recalled the position of three human characters relative to me. I confirmed my facing direction around them, and then I chose the target location. | 100% |
|  | I recalled what I experienced during walking period. | 21% |

## Notes

**Conflict of interest statement**: no competing financial interests

